# Interstrain Recombinants of Human Cytomegalovirus Reveal Complex Genetic Correlates and Epistasis Influencing Glycoprotein Display, Virion Infectivity and Spread Characteristics

**DOI:** 10.1101/2025.09.22.677782

**Authors:** Christopher Peterson, Ian T. Bailey, Jean-Marc Lanchy, Ivan Gallego, Brent J. Ryckman

## Abstract

Most of the nucleotide diversity in the human cytomegalovirus (HCMV) genome is due to approximately 17 genes with 2-14 alleles each. These allelic genes are interspersed among longer stretches of highly conserved sequences with signatures of extensive recombination that would shuffle the allelic genes into a vast number of allelic haplotypes. Bacterial artificial chromosome clones derived from 3 independent clinical isolates (TB40/e (TB), TR and Merlin (ME)) display dramatic differences in the abundance of entry-mediating glycoproteins gH/gL/gO and gH/gL/UL128-131, virion infectivity and efficiency of cell-free and cell-to-cell modes of spread. Of these, TB and ME are the most phenotypically different and share only 2 of the 17 allelic genes. A set of recombinant HCMV was generated by coinfecting cells with TB and ME and restriction fragment length polymorphism (RFLP) analyses demonstrated complex crossover patterns. Most recombinants were either “TB-like” with much more gH/gL/gO than gH/gL/UL128-131, or “ME-like” with much more gH/gL/UL128-131. This correlated with a TB or ME UL128 sequence, consistent with a G/T polymorphism affecting UL128 pre-mRNA splicing. One recombinant had a gH/gL/gO:gH/gL/UL128-131 ratio of 0.8, suggesting genetic determinants beyond UL128. Virion infectivity correlated with TB versus ME-like glycoprotein display, but intragroup variability indicated additional factors and variability in spread efficiency and the contribution of cell-free and cell-to-cell spread modes indicated an influence of characteristics beyond virion infectivity. Results suggest that the relationships among these three phenotypes are not strictly causal and that all three phenotypes are genetically complex and influenced by epistasis among polymorphic loci across the genome.

**IMPORTANCE:** The emerging picture of the genetic diversity of HCMV *in vivo* prompts a reevaluation of how *in vitro* characterized viral phenotypes, such as the abundance of the various glycoproteins displayed on the virion envelope and tendency towards cell-free or direct cell-to-cell spread, reflect viral characteristics *in vivo*. A small sampling of the *in vivo* genetic diversity of HCMV has been examined in the laboratory and the genetic correlates of *in vitro* phenotypes are unclear. Engineered mutations can yield different phenotypic effects when introduced on to different genetic backgrounds, suggesting complex epistasis among polymorphic loci across the genome. By generating a set of recombinants derived from two phenotypically and genotypically distinct parental clones, this work offers an approach to characterize complex genetic correlates of viral phenotypes, which in turn may enhance efforts to link genomics data with *in vivo* viral behavior and clinical outcomes.

## INTRODUCTION

One of the 9 human herpesviruses, human cytomegalovirus (HCMV) has an average global prevalence rate of 83%, although this varies by region and stratifies by age, as the longer an individual lives the more chances to seroconvert (1). Infections are typically mild in immunocompetent individuals but can be dangerous in those with compromised or naive immune systems, including those living with HIV/AIDS, transplant recipients, and newborns (2–5). At least 1 in 5 infants are born each year in the United States with HCMV infection, leading to problems ranging from stunted growth, to hearing loss, and even microcephaly. Previous infection does not offer broad protection against subsequent infections or prevent intrauterine infection of developing fetuses (6–8). Consequently, congenital HCMV infection is more common in regions of the world with higher seroprevalence among adults (9). Success of interventions including vaccines and drug therapeutics have been limited (10–12), and this may be due in part to an inadequate understanding of how the genetic variation of HCMV impacts viral phenotypes.

The scope and dynamics of HCMV genetic variation are becoming more clear (13–23). The majority of nucleotide (*nt)* diversity in the HCMV genome is clustered into discrete regions interspersed among longer stretches of highly conserved sequences. High linkage-disequilibrium (LD) in the variable regions indicates that the genes present are stable, fixed alleles and low LD in the conserved sequences suggests recombination that would shuffle the variable, allelic genes into a vast number of combinations or “allelic haplotypes”. The intrahost *nt* diversity of HCMV can be comparable to that of some RNA viruses, likely reflecting mixed haplotype infections rather than rapid genetic drift, more consistent with the higher fidelity of DNA replication (24). *In vitro* characterization of HCMV phenotypes have involved a limited sampling of the apparent *in vivo* genetic diversity and questions about artifactual *in vitro* genetic drift are complicated by a lack of understanding of the genetic correlates of phenotypes and the influence of epistasis among the allelic genes across the genome.

We’ve reported extensive phenotypic variations among 3 HCMV BAC clones derived from distinct clinical isolates, TB40-BAC4 (TB), TR-BAC (TR), and Merlin-BAC (ME). These 3 clones differ in the amounts of 2 entry-mediating glycoprotein complexes in the virion envelope, gH/gL/gO (“trimer”) and gH/gL/UL128-131 (“penatmer”) (25, 26). The shared gH/gL portion of these complexes contributes to the fusion apparatus, acting as a cofactor for the fusion protein gB, whereas the accessory subunits gO and the UL128-131 subcomplex are receptor-binding domains that influence cell-type specific tropism (27–30). TB and TR virions contain large amounts of trimer and low levels of pentamer, often below the level of detection by immunoblot, whereas ME expresses less total gH/gL, the bulk of which is pentamer (25, 26). The ME BAC clone was engineered with tetracycline (tet)-operator sequences in the UL131 promoter such that progeny virus produced in cells expressing the tet-repressor protein (TetR) are drastically reduced in pentamer and slightly increased in trimer (25, 31). The virion amounts of trimer and pentamer correlate with the infectivity on different cell types and the known receptors. Trimer binds PDGFRα and facilitates entry to fibroblasts (30). Consistent with this, TB and TR are considerably more infectious than ME on fibroblasts, and the infectivity of ME is greatly increase by propagation in TetR cells (25). Pentamer promotes infection of epithelial cells by binding NRP-2 and OR14I1(29, 32). While the infectivity of all 3 clones is lower on epithelial cells, the comparisons between clones are generally similar indicating that the low levels of pentamer in TB and TR virions are sufficient and trimer plays an important role for infection of epithelial cells as well as fibroblasts (25, 33). This, coupled with the low (or lack of) PDFGRα expression in these epithelial cells, suggests other trimer receptors (32, 34–36). The relatively high virion specific infectivity of TB and TR is consistent with the ability of these strains to spread more efficiently via diffusion of progeny released to the culture supernatants (i.e. “cell-free spread”), whereas ME is more restricted to direct cell-to-cell spread, consistent with low virion infectivity (37).

The precise genetic basis of phenotypic differences among TB, TR and ME are unclear. A G/T single *nt* polymorphism (SNP) between TB and ME was suggested to affect a predicted splice site on the UL128 pre-mRNA, contributing to the low pentamer observed in TB virions (38). There are likely other factors since TR is also low in pentamer but is like ME at this G/T SNP residue. Indeed, Zhang et al demonstrated that in addition to high expression of the UL128-131 proteins, ME is also low in gO expression compared to TR, and this low gO expression might be related to a reduced expression of UL148, an ER resident protein that regulates endoplasmic reticulum-associated degradation (ERAD) of gO (39, 40). The virion amounts of pentamer and trimer influence specific infectivity, but there are likely other important factors since 1) infectivity of ME virions produced in TetR cells is comparable to that of TR despite containing far less trimer and the infectivity of TB is far greater than that of TR despite comparable levels of trimer (25), and 2) allelic variation in gO can impact virion infectivity and the kinetics of membrane fusion regulated by gH/gL/gO binding to the receptor PDGFRα (41, 42). Likewise, proclivity for cell-free and cell-to-cell spread is likely impacted by factors beyond virion specific infectivity since 1) despite producing highly infectious intracellular virions, spread of TB was highly sensitive to inhibition by neutralizing Abs (nAb), 2) the rate of cell-to-cell spread by ME was comparable to that of the cell-free spread of TB, and 3) spread of ME in TetR cells was much more resistant to nAb inhibition than that of TB despite the enhanced extracellular virion infectivity (37). Thus, the respective specializations of TB and ME to cell-free and cell-to-cell-spread are not merely reflections of virion infectivity. Rather, ME seems capable of a specific mechanism(s) that promotes cell-to-cell spread that TB is not. Together, these observations suggest that each of the 3 characterized phenotypes, 1) trimer/pentamer display, 2) virion specific infectivity, and 3) spread characteristics are complex, polygenic phenotypes influenced by genome-wide epistasis. Here we describe the generation of set of TB-ME recombinants that display a wide range of intermediate phenotypes and disrupt the apparent correlations between them.

## RESULTS

### Construction of TB-ME recombinant library

Most of the *nt* diversity within the HCMV genome is clustered into discrete regions distributed throughout otherwise highly conserved sequences (20). High LD indicates that the genes in these variability clusters are distinct, fixed alleles. Seventeen of these genes were characterized to have between 2 and 14 alleles (17) (Fig 1). Table 1 depicts the “allelic relatedness” of the three reference clones used in our previous studies, TB TR and ME. TR shares a different set of 4 alleles with both TB and ME whereas the 2 most phenotypically distinct clones, TB and ME, share only 2 alleles. To define the genetic correlates and genome-wide-epistasis influencing and connecting the 3 phenotypes, 1) trimer:pentamer ratio, 2) virion infectivity and 3) spread mode preference, a set of inter-clone recombinants was made with TB and ME as the parents.

**Figure 1.**
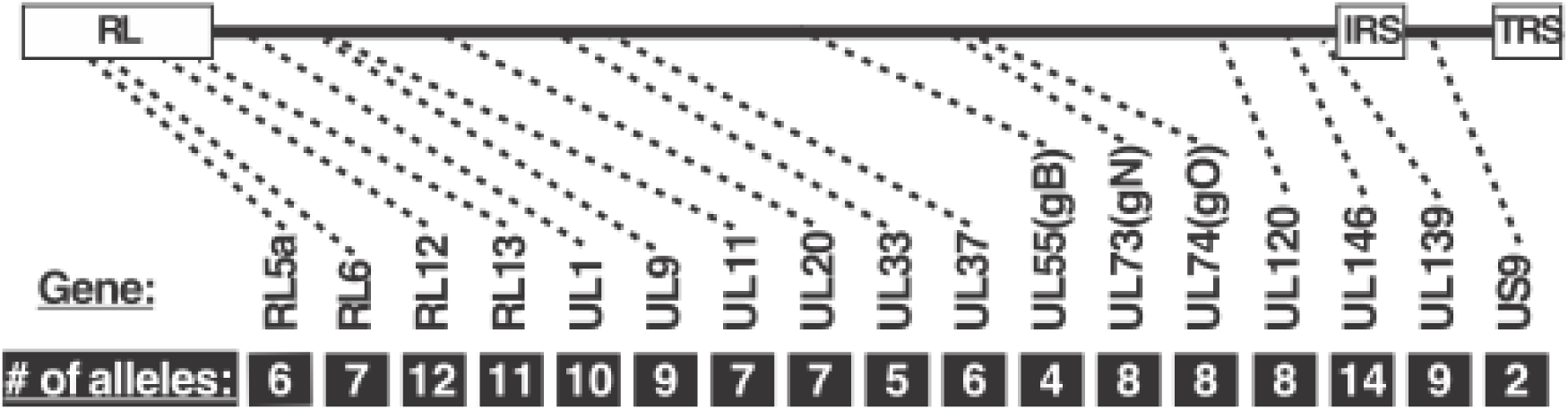
The Allelic Diversity of HCMV. A diagram of the HCMV genome indicating the approximate location of allelic genes and number of alleles each based on the analyses of Lassalle and Suarez (17, 20).

**Table 1.**
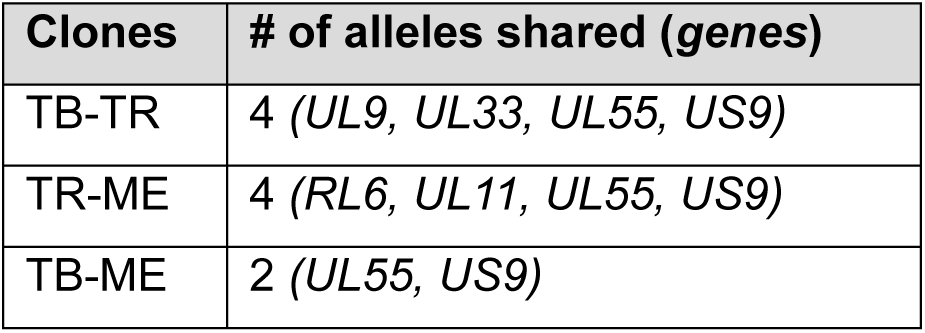
Allelic relatedness of HCMV BAC clones.

| **Clones** | **# of alleles shared (*genes*)** |
| --- | --- |
| TB-TR | 4 (*UL9, UL33, UL55, US9*) |
| TR-ME | 4 (*RL6, UL11, UL55, US9*) |
| TB-ME | 2 (*UL55, US9*) |

Parental TB and ME containing mCherry (mC) or GFP genes in place of US11 (37) were used to coinfect cultures of HFFtet cells (Fig 1). Note that parental ME-GFP was propagated on TetR-expressing human foreskin fibroblasts (HFFtet), suppressing the expression of UL131 to enhance the virion infectivity and reduce selection against gH/gL/UL128-131 (“MT-GFP”) (25, 31). Cells infected by both parental viruses (i.e., GPF+/mC+ cells) were batch collected by FACS, replated and maintained for 8 days at which time progeny virus was harvested from the culture supernatant and infectious units (IU) quantitated by fluorescent marker; 1 x 10^6^ (GFP) and 6 x 10^6^ (mC) infectious units (IU) per mL. Since no new uninfected cells were added, 100% of the progeny virus were derived from cells coinfected by TB and ME, thus potential recombinants. Moreover, since the GFP and mC marker genes were inserted into the homologous locus in both parental genomes (i.e., US11), no recombinant progeny could harbor both markers without major, and likely highly deleterious genetic abnormalities. Supernatant virus derived from dual GFP+/mC+ cells was passed through a 0.4 um filter to remove virion aggregates and detached, infected cells and then used to infect HFFtet cells at a low multiplicity to disfavor infection by more than 1 virion per cell. After 2 days, cells were FACS sorted to collect single GFP+ and mC+ infectious centers into 96 well plates along with fresh, uninfected HFFtet cells. The use of HFFtet cells throughout this process reduced selection against any “ME-like” viruses, high in gH/gL/UL128-131 (31).

Focal expansion from infectious centers was monitored visually for 10-20 days. Approximately 70% of wells showed single GFP or mC focal expansion and these were harvested for analysis. The remaining 30% of wells either had both GFP and mC or showed no focal expansion and were discarded. Whereas both parental viruses formed large, dispersed foci on HFFtet cells, indicative of efficient cell-free spread, the putative recombinant isolates showed a range of focal patterns from large and dispersed to small and compact (Fig. 2F). Fifteen isolates showing varied focal patterns were selected for expansion and further characterization. Two were subsequently discarded during characterization of recombinant genome structure (see below).

**Figure 2.**
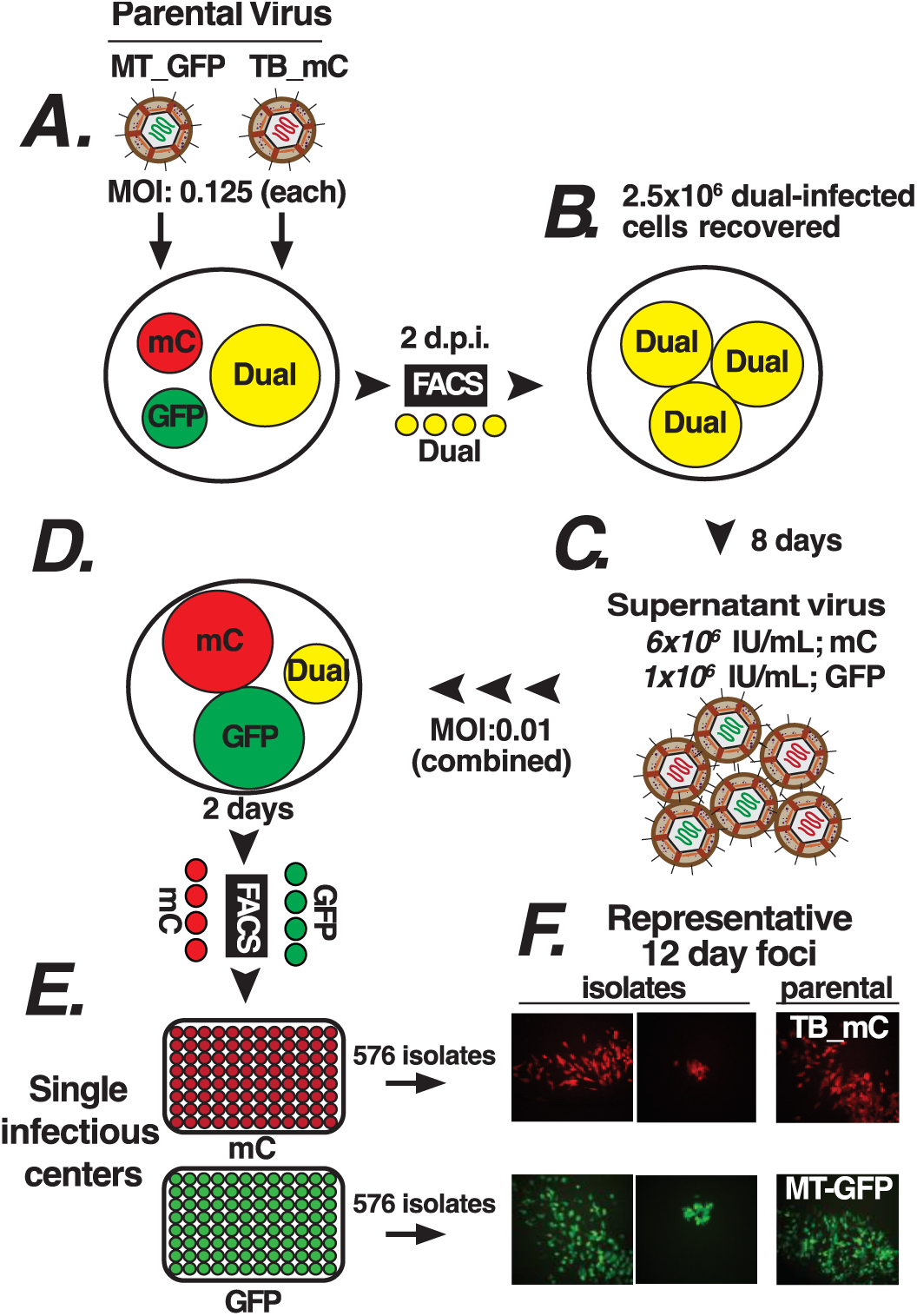
Isolation of virus progeny from TB-ME coinfected cells. (A) HFFtet cells were coinfected with parental TB-mCherry (mC) and MT (ME produced in HFFtet cells)-GFP (B) Dual-infected GFP+/mC+ cells were collected by FACS at 2 days post infection (dpi). (C) Progeny virus was purified after 8 days and GFP and mC infectious units (IU) determined. (D) Fresh HFFtet cells inoculated at low multiplicity, and (E) single GFP+ and mC+ cells collected at 2 dpi into 96 well plates of HFFtet cells as infectious centers. (F) Isolates recovered displayed a range of foci phenotypes.

### Restriction fragment length polymorphism (RFLP) analysis revealed complex recombination patterns

Recombinant genome structure was assessed by PCR-RFLP analysis. A set of PCR primers was designed with perfect conservation to both parental virus sequences but where sequence polymorphisms near the middle of the PCR amplicon disrupted a restriction nuclease site in one of the two parents (Table 2). Figure 3A shows an example for a predicted PCR amplicon within UL74. The 790 base pair (bp) amplicon from the ME sequence contained an Nde I site near the middle resulting in fragments of 422 and 368 bp, which comigrated during electrophoresis. T>G and G>A polymorphisms in TB (Table 2) prevented Nde I digestion. In this experiment, the PCR input template was titrated from 100% TB to 100% ME demonstrating detection of an approximately 95:5 mixture of parental viruses. Figure 3B shows an example analysis of a TB-ME recombinant isolate denoted E7 compared to both parental viruses. The UL74 amplicon of E7 was resistant to Nde I, like the parental TB, whereas the UL98 amplicon was sensitive to Sac I, like the parental ME. This indicated that isolate E7 was a TB-ME recombinant with least one recombination crossover point between UL74 and UL98. All 15 isolates were analyzed at 13 loci across the genome and each locus was assigned as “TB” or “ME” (Fig. 3C). An additional locus, UL128, was assigned as “TB” or “ME” by Sanger sequencing. As mentioned above, 2 of the 15 isolates were discarded due to discrepancies in assigning TB vs ME of at least one RFLP locus in 3 independent analyses, suggesting that these isolates were of mixed genotype. Of the remaining 13 isolates, 10 were confirmed recombinant, showing a variety of complex crossover patterns. The RFLP patterns of 2 isolates matched the parental TB (E9 and F7) and 1 matched the parental ME (G6). However, given that there were 10’s of kbp between RFLP loci, it is not possible to conclude that these 3 isolates were *bona fide* “non-recombinant parental.” There may have been crossover site between RFLP loci. Indeed, these isolates showed phenotypic differences compared to the parental viruses (see below).

**Figure 3.**
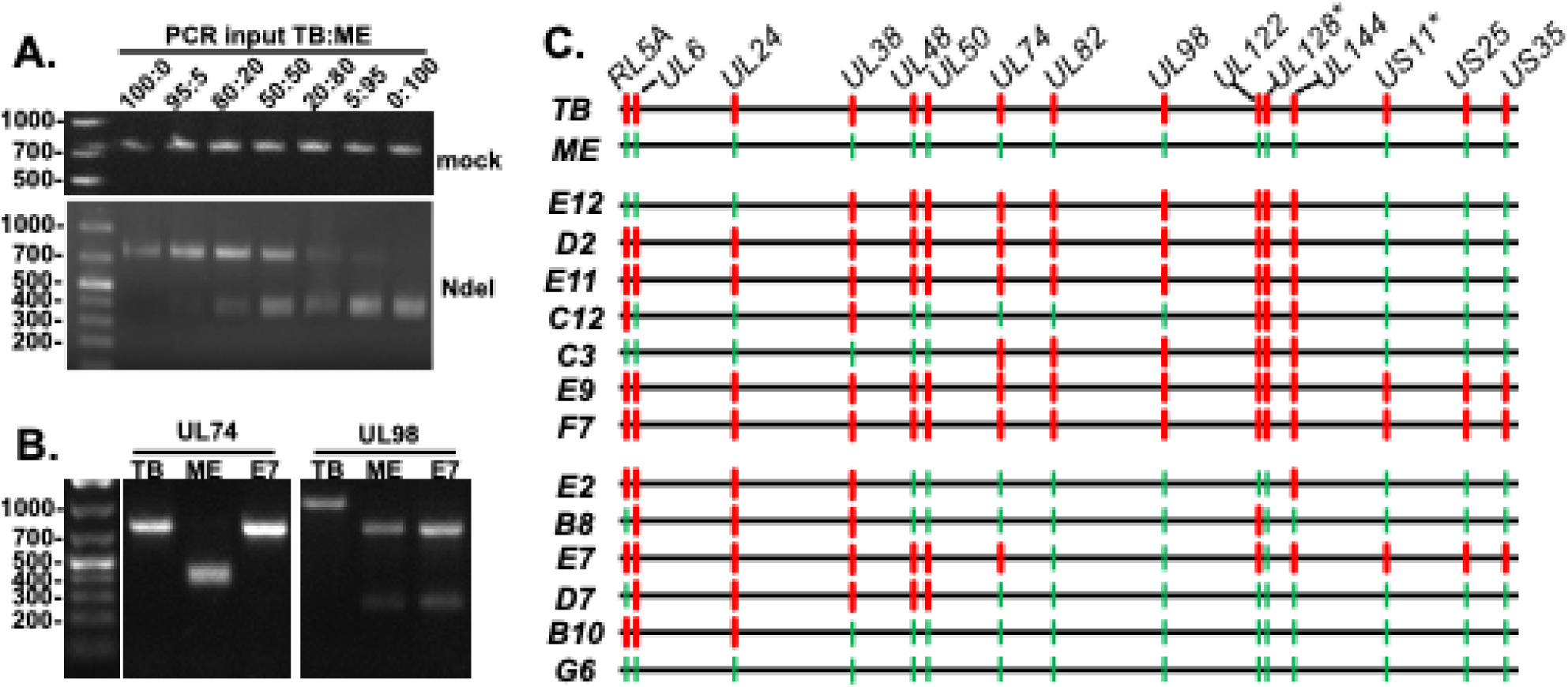
Restriction fragment length polymorphism (RFLP) analysis of recombinant genome structure of TB/ME coinfection progeny isolates. (A) TB and ME BAC DNA was mixed in the indicated ratio as input template for PCR reactions with primers to UL74 sequences (Table 2). PCR amplicon was mock-digested (top) or digested with Nde I (bottom) and then analyzed by agarose gel electrophoresis and stained with ethidium bromide. (B) DNA extracts from cell infected with parental TB, ME or the isolate E7 were PCR amplified with primers within the UL74 or UL98 loci, then digested with Nde I or Sac I, respectively and analyzed by agarose gel electrophoresis and stained with ethidium bromide. (C) TB versus ME assignments of loci based on RFLP as in Table 2. * Assignment of UL128 based on sequencing, not RFLP, and US11 locus based on visible GFP or mC fluorescence.

**Table 2.**
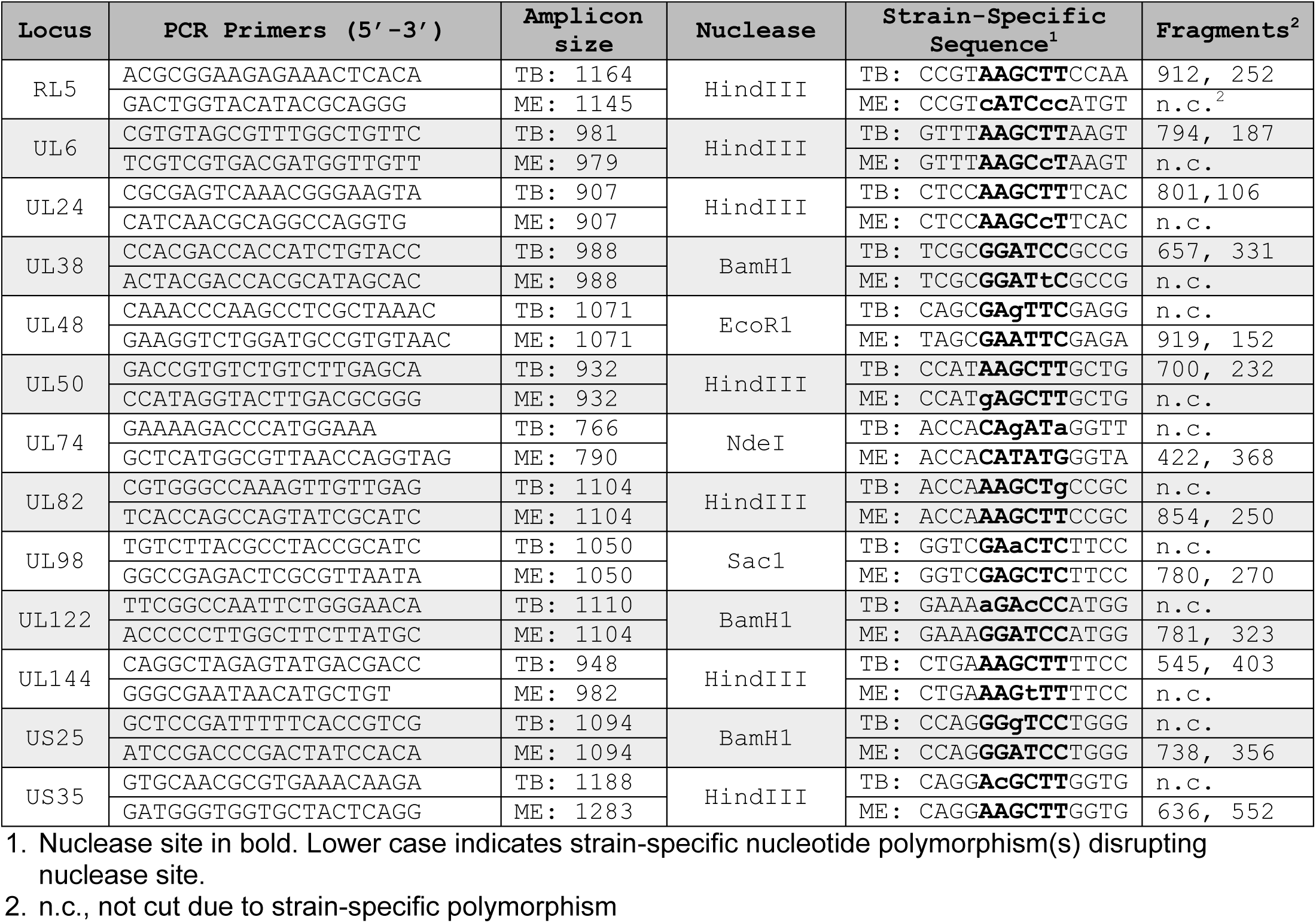
Predicted PCR amplicon sizes and restriction fragment patterns.

### Phenotypic analysis of recombinant HCMV

While recombinants were isolated and expanded in HFFtet cells to reduce selection against the UL128-131 locus for “ME-like” viruses, high multiplicity (i.e., single-round infection) “burst” stocks were generated on HFF lacking TetR expression to analyze phenotypes.

1. *Display of gH/gL/gO (trimer) and gH/gL/UL128-131 (pentamer).* The relative abundance of gH/gL/gO and gH/gL/UL128-131 (i.e., trimer:pentamer ratio) of the recombinant isolates was analyzed by immunoblot and band densitometry. Consistent with previous results (26), TB virions contained far more gH/gL/gO than gH/gL/UL128-131 with a ratio of 3.9 in the experiment shown, whereas ME virions contained more gH/gL/UL128-131 than gH/gL/gO with a ratio of 0.4 (Fig 4A and B). The ratios of most recombinants were either much greater than 1.0, like the parental TB (Fig. 4A) or much less than 1.0, like the parental ME (Fig 4B). Only one isolate (B8) had a nearly balanced amount of gH/gL/gO and gH/gL/UL128-131 with a ratio of 0.8. Given this ratio was lower than 1.0, isolate B8 was included among the “ME-like” set for further analyses.
2. *Virion specific infectivity.* We previously reported that the per genome infectivity of TB stocks were log-folds higher than for ME stocks (25, 37, 43). The specific infectivity of the recombinant isolates corresponded categorically to the trimer:pentamer ratios with “TB-like” recombinants displaying much higher infectivity than the “ME-like” viruses (Fig. 5A and B). Among the TB-like recombinants, E12, D2, E11, E9 and F7 were marginally less infectious than the parental TB (1.5 to 4-fold) whereas C12 and C3 were 10.2 and 26.3-fold less infectious, respectively (Fig. 5A). Among the ME-like recombinants, E2 was 15.4-fold more infectious than the parental ME, whereas B8, E7, and D7 were not statistically different from ME. Recombinants B10 and G6 were 14.8- and 31.7-fold less infectious than ME, respectively. However, these differences were not statistically significant likely due to variance reflecting the lower limits of the infectivity measurement.
3. *Spread characteristics.* Despite the dramatic differences between TB and ME in extracellular virion infectivity, we previously showed that the clones spread through HFF cultures at comparable rates, TB predominantly via diffusion of highly infectious extracellular virions that can be efficiently blocked by nAb and ME predominately via direct cell-to-cell spread that is less sensitive to nAb inhibition (37). There was considerable variation in spread efficiencies among the TB-like recombinants (Fig. 6A). Recombinants D2, E11, E12, C12 and C3 each spread more efficiently than the parental TB (2.8, 4.2, 2.1, 1.8, and 4.6-fold, respectively), whereas spread of E9 was 4.6-fold reduced. There was less variation among the ME-like recombinants with only D7, B10 and G6 statistically different from the parental ME (1.3, 1.5 and 1.7-fold greater, respectively) (Fig. 6B). The reduced spread of E9 compared to parental TB correlated with a difference in specific infectivity, (i.e., approximately 4-fold reduced in both specific infectivity and spread). In contrast, none of the enhanced spread could be explained by enhanced virion specific infectivity as all these recombinants had decreased infectivity compared to the parental TB or ME (compare Figs. 5A,B with 6A,B).

**Figure 4.**
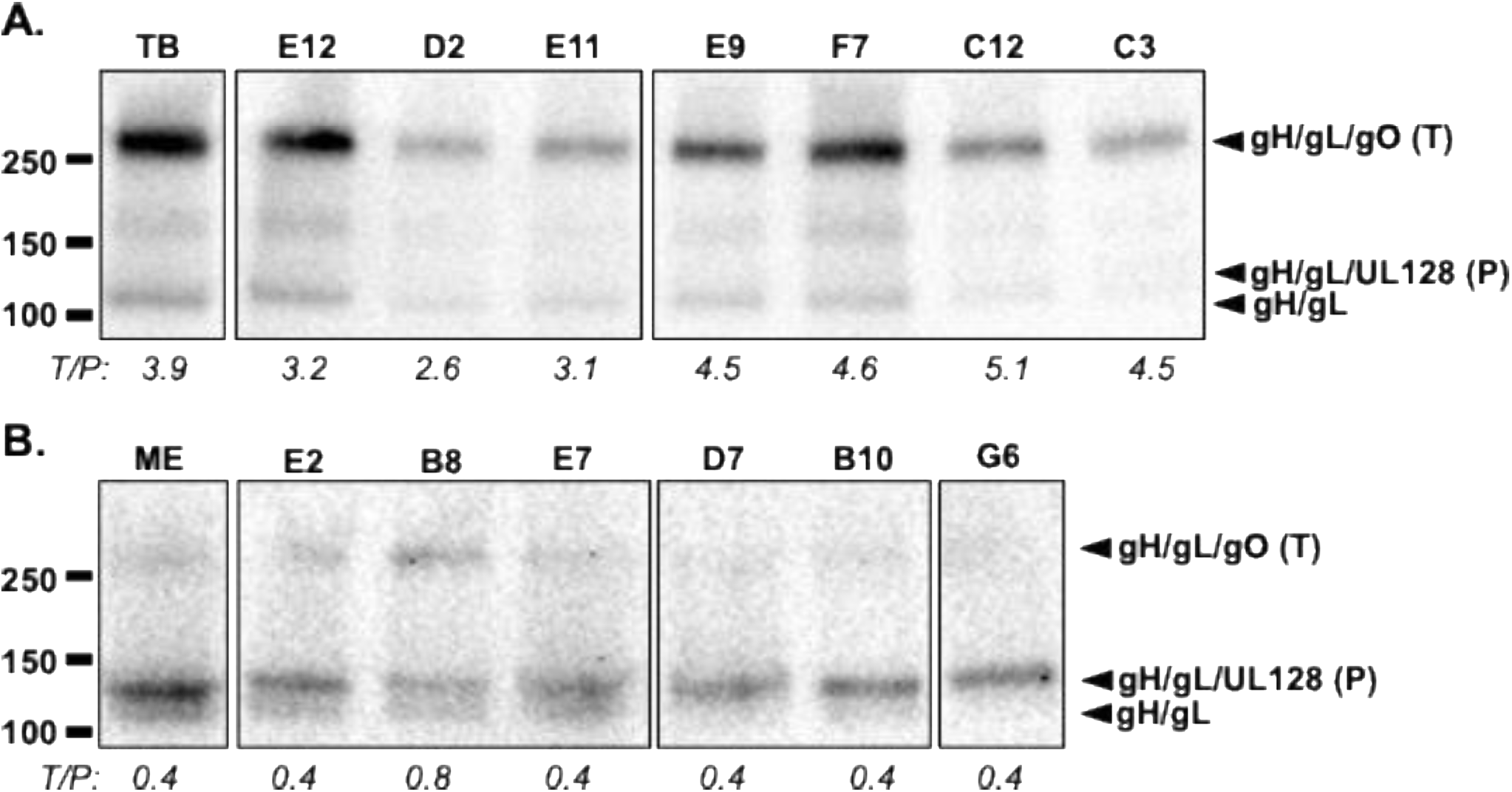
Immunoblot analysis of gH/gL complexes in parental and recombinant HCMV. Extracts of HCMV virions pelleted from HFF culture supernatants were separated on non-reducing 4-20% gradient SDS-PAGE gels, transferred to nylon membranes and probed with anti-gL antibodies. Mobility markers indicated in kDa (left). Locations of gH/gL/gO (trimer “T”), gH/gL/UL128 (pentamer “P”), and gH/gL indicated to the right of each panel. Loads were not normalized to number of virions. Ratio of band densities trimer/pentamer (T/P) indicated beneath each lane in italics as determined using FIJI V.2.16/1.54p. All lanes shown were digitally cropped from the same gel images in A and B, respectively, and are representative of at least 3 biological replicates.

**Figure 5.**
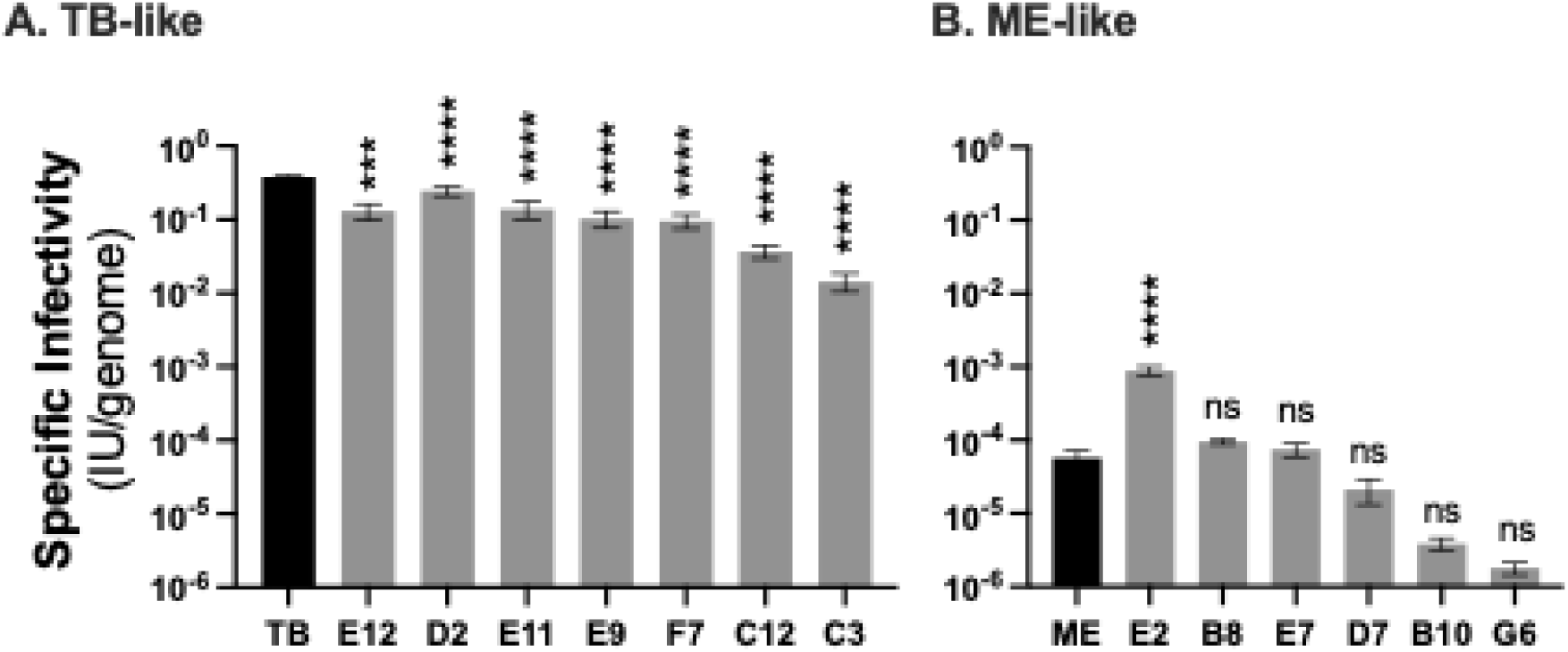
Specific infectivity of parental and recombinant HCMV. Extracellular HCMV stocks were quantified by qPCR for viral genomes and infectious units (IU) determined by flow cytometry quantification of GFP or mC expression in HFF cells 3 days post infection. Average IU/genome from at least 3 biological replicates are plotted with error bars representing standard deviation. Asterisks indicate *P* values of differences between recombinants and the parental TB (A) or ME(B), *** < 0.001. **** <0.0001, ns = not significant calculated by one-way analysis of variance (ANOVA) with Dunnett’s post hoc comparing each recombinant to TB (A) or ME (B).

**Figure 6.**
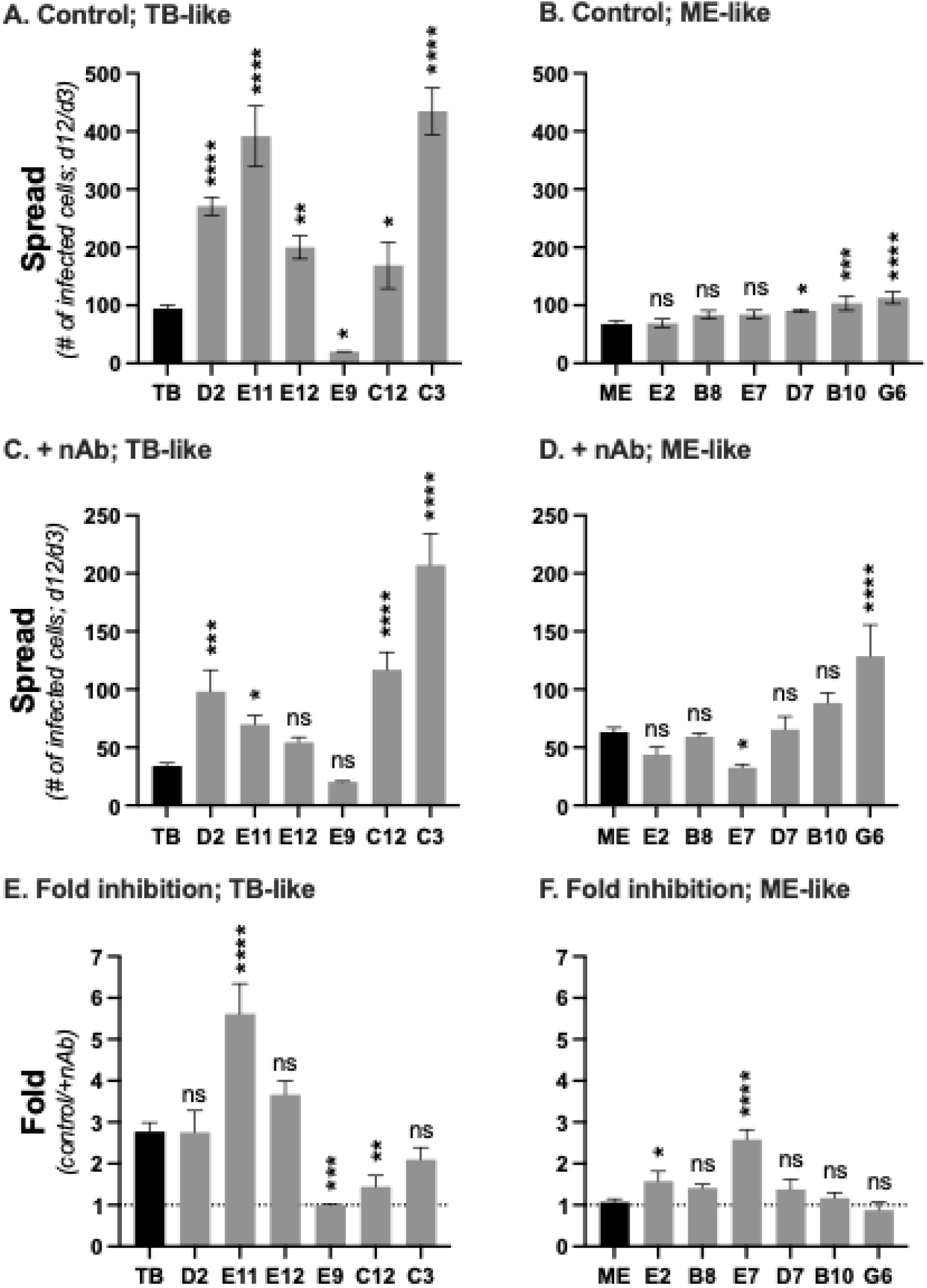
Spread characteristics of parental and recombinant HCMV. (A and B). Confluent monolayers of HFF cells were infected at low MOI with the parental (TB or ME) or recombinant isolates. At 3 and 12 days postinfection, the number of infected cells was determined by flow cytometry detection of GFP or mC. Plotted are the average number of infected cells at day 12 per infected cell at day 3 in 3 biological replicate experiments. Error bars indicate the standard deviation. (C and D). As in A and B except cultures were maintained in medium supplemented with 50 μg/mL anti-gH mAb 14-4b. (E and F). The ratio of values A:C and B:D, error propagated. Asterisks indicate *P* values of differences between recombinants and the parental TB (A, C, E) or ME (B, D, F). *\* ≥* 0.05, ** < 0.01, *** < 0.001. **** <0.0001, ns = not significant calculated by one-way analysis of variance (ANOVA) with Dunnett’s post hoc comparing each recombinant to TB (A, C, E) or ME (B, D, F).

In parallel spread experiments, nAb were included in the culture medium to assess the contribution of extracellular virus to spread (Figs 6C-F). Consistent with the previous results (37), spread of parental TB was strongly inhibited by nAb, indicative of predominantly cell-free spread, whereas spread of ME was not affected by nAb, indicative of predominantly direct cell-to-cell spread. Spread of recombinant E11 was 2-fold more sensitive to nAb than the parental TB suggesting that the enhanced spread efficiency of this recombinant was due to increased cell-free spread. In contrast, recombinant C12 was 1.9-fold less sensitive to nAb inhibition, consistent with an increased capacity for direct cell-to-cell spread. Spread of recombinant E9, which was 4.6- fold reduced compared to TB (Fig. 6A), was totally unaffected by the presence of nAb (Fig 6C,E), further suggesting that the reduced spread efficiency was due to decreased infectivity of the extracellular virions and the remainder of spread was predominantly via a low efficiency cell-to-cell mechanism. Among the ME-like recombinants, spread of both E2 and E7, which were comparable to the parental ME in efficiency, were more sensitive to nAb inhibition (1.5- and 2.5-fold respectively) indicating more reliance on cell-free spread.

## DISCUSSION

The unveiling of the vast and complex nature of HCMV genetic diversity compels reevaluation of how *in vitro* characterizations of HCMV phenotypes reflect viral characteristics *in vivo*. For example, the tendency of clinical isolates to display a cell-associated phenotype in culture is has been viewed to support the notion that HCMV *in vivo* is principally cell-associated and that a cell-free phenotype is an *in* vitro artifact (44). Indeed, leukocyte depletion has been linked to reduced transmission of HCMV during blood transfusions, arguing against large amounts of infectious, cell-free HCMV in the blood (45, 46). While this is consistent with a model of hematogenous dissemination of HCMV via monocyte/macrophages (47, 48), these were small scale studies that do not offer broad insights into the roles of cell-free and cell-associated virus in other aspects of HCMV pathogenesis. Moreover, while cell-association is a commonly observed phenotype for clinical isolates, cell-free phenotypes during initial culturing have also been reported (18, 49). Comprehensive understanding of the *in vivo* role of phenotypes like cell-free and cell-to-cell spread will benefit from a more complete understanding of the associated genetic signatures.

The HCMV BAC clones TB and ME display opposite extreme *in vitro* phenotypes including the levels of the entry-mediating glycoprotein complexes gH/gL/gO and gH/gL/UL128-131, virion specific infectivity and preference for cell-free or cell-to-cell modes of spread (25, 26, 37, 40, 41, 43). Consistent with phenotypic extremes, TB and ME are mismatched at 15 of 17 allelic genes. To study how genome-wide epistasis might influence phenotypes, we generated a set of TB-ME recombinants by isolating progeny from cells coinfected by TB and ME and assessing recombinant genome structure by targeted PCR-RFLP. One limitation to this approach was the possibility that stochastic PCR amplification of one of multiple homologous targets over the others in the early cycles could give the appearance of genetic purity when in fact the isolate was of mixed genotype. This was addressed with a TB-ME genome titration experiment that demonstrated the ability to detect approximately a 5% minor variant, and a stringent standard was set of agreement among 3 independent experiments across all RFLP loci for each isolate. Indeed 2 of the original 15 isolates selected for analysis were discarded due to inconsistent results in RFLP patterns in the 3 experiments. Thus, while the PCR-RFLP may be considered low resolution in not showing exact locations of crossovers, the approach demonstrated that the isolates were *bona fide* TB-ME recombinants with a variety of complex crossover patterns.

The first major phenotypic discrepancy between TB and ME that we reported was that TB virions contained far more gH/gL/gO than gH/gL/UL128-131, whereas ME virions contained more gH/gL/UL128-131(26). The genetic basis of this difference appears to be complex. A G-to-T polymorphism in TB relative to ME was suggested to reduce the splicing efficiency of the TB UL128 pre-mRNA and when engineered into ME, reduced the assembly and expression of gH/gL/UL128-131 into ME virions (38, 50). More recently, the clinical isolate TB40/E (the source of the TB40-BAC4 clone that we abbreviate as “TB”) was shown to contain both variants and the effect of the G/T polymorphism on UL128 abundance was corroborated (51). Our results align with these reports as most of the TB-ME recombinants analyzed retained either a “TB-like” or “ME-like” trimer:pentamer ratio and this correlated with the presence of the TB and ME UL128 sequence. However, UL128 polymorphism is unlike to be the sole factor dictating trimer:pentamer levels since, 1) TetR repression of pentamer expression in ME does not result in a compensating increase in virion levels of gH/gL/gO (25), and 2) the TR BAC clone is like ME at this UL128 intron splice site, but is trimer-high/pentamer-low like TB (25). Zhang et al reported that in addition to high expression of the UL128-131 genes, ME is particularly low in gO expression and this might be due to low expression of UL148, which can protect gO from ERAD (39, 40). This may help explain why the trimer:prentamer ratios among the TB-like recombinants were more variable then among the ME-like recombinants, i.e., there was more dynamic range of gO expression. More recently, Weiler et al. suggested that both the TB and ME trimer:pentamer ratio extreme phenotypes might be anomalous based on their observation that 8/8 recent clinical isolates analyzed had balanced trimer:pentamer ratios (52). Indeed, one of our recombinants (B8) showed a nearly balanced trimer:pentamer ratio, suggesting that it is possible for a single HCMV genome to contain the right collection of genetic polymorphisms to epistatcially result in a balanced trimer:pentamer ratio. However, it still may be premature to suggest that a balanced trimer:pentamer ratio is more representative of *in vivo* HCMV than the extremes of TB and ME. Weiler et al. did not provide data to address the possibility of the isolates being genetic mixtures of TB- and ME-like variants, which would be expected to yield balanced trimer:pentamer ratios on immunoblots. Moreover, the clinical isolates analyzed were all collected in the same location within a relatively short period of time, decades after the clinical collection of the TB and ME isolates in distinct locations (49, 53). Our observations with TB:ME recombinants support the idea that trimer:pentamer ratio of HCMV is controlled epistatically by variation in multiple loci across the genome and that whether a clinical isolate presents with balanced or imbalanced trimer:pentamer ratio may be due to the dominance of specific haplotypes circulating in that particular geographic location and time.

The relationship between the extreme disparity in trimer:pentamer ratio between TB and ME and their extreme difference in specific infectivity is unclear. TB virions are highly infectious whereas the infectivity of ME approaches the lower limits of our infectivity assay (25, 37, 41). The ME clone used in our studies contains Tet operator sequences in the UL131 promoter such that propagation in TetR-expressing fibroblasts results in virions with dramatically less pentamer and slightly more trimer (we denoted these virions “MT”) (25, 31, 38). Whether the enhanced extracellular infectivity of MT virions compared to ME virions was the result of slightly more gH/gL/gO or the dramatic reduction in gH/gL/UL128-131 was not clear. Note that while attachment and entry events are critical to infectivity, the readout of infectivity in these experiments was viral gene expression, many steps downstream of virus entry. A recent study suggested that the pentamer complex can induce cellular signaling pathways during entry into fibroblasts that can subsequently inhibit IE gene expression, thus reducing apparent infectivity (54). This idea is consistent with our observation that the specific infectivity of the TB:ME recombinants grouped categorically with the “TB- or ME-like” trimer:pentamer ratio status. Moreover, this offers a potential explanation for the apparent instability of the UL128-131 genes of the ME clone and their relative stability in the TB and TR clones (50). The 15.4-fold enhanced infectivity of recombinant E2 compared to parental ME indicates that a pentamer-induced inhibition might be least partially compensated by increases in other mechanisms leading to viral gene expression such as nuclear docking, uncoating or subversion of cellular antiviral state mechanisms. Conversely, all of the TB-like recombinants were less infectious than the parental TB (1.5- to 26.3-fold), suggesting that the recombinatorial mixing of TB and ME genes can reduce the efficiency of one or more mechanisms during the establishment of infection, consistent with the observation that the TB clone is highly specialized to cell-free spread (37). Together, these observations highlight the complex, multi-mechanism nature of “infectivity” using viral gene expression as the readout.

In our usage, the phenotype “spread” represents the totality of mechanisms involved in the viral replication cycle over multiple generations, and again TB and ME show dramatic differences. TB and ME spread at similar rates in fibroblasts cells (i.e., the number of new infected cells per initially infected cell, over time) but whereas spread of TB relies predominately on progeny virions released to the culture supernatant that can be efficiently blocked by nAb (i.e., “cell-free” spread), ME predominately spreads directly cell-to-cell and this is much less sensitive to nAb inhibition (37). There is a logical connection between the production of highly infectious extracellular virions and efficient cell-free spread, but the mechanistic basis for efficient cell-to-cell spread is more complicated. While repression of pentamer during ME replication dramatically improves extracellular virion infectivity, this does not come at the expense of efficient cell-to-cell spread (37). Moreover, the intracellular virions of TB are far more infectious than those of ME when liberated from the infected cells by sonication, yet cell-to-cell spreads of TB is comparatively inefficiency (37). Together these observations indicate that some aspect(s) of ME physiology allows for efficient cell-to-cell spread despite poorly infectious progeny virions, and TB is deficient, despite producing highly infectious intracellular virions. In our studies there was more variability in spread characteristics among the TB-like recombinants than among the ME-like set. The increased spread of recombinants D2, E11, E12, and C3 compared to parental TB was accompanied by either equal or greater sensitivity to nAb inhibition, suggesting predominat cell-free spread. Given that these recombinants were reduced in specific infectivity compared to TB, it may be that the mixing of TB and ME genes resulted in the more efficient production and release of greater quantities of progeny to the culture supernatant. In contrast, spread of isolate E9 was reduced relative to TB, which might be explained by this isolate’s reduced specific infectivity. Conversely, spread of recombinant C12 was both more efficient and less sensitive to nAb inhibition than TB, suggesting the acquisition of ME genes that allowed for more efficient cell-to-cell spread. Among the ME-like recombinants, E2 and E7 stand out as spreading comparably to ME, but with greater sensitivity to nAb inhibition. For E2, this was consistent with an increased specific infectivity of extracellular virions, suggesting a shift towards a greater contribution of cell-free spread. However, the increased nAb inhibition of E7 could not be explained by infectivity. The observation that recombinatorial mixing of TB and ME genes resulted in a variety of different spread characteristics underscores that “spread” as measured here is a complex, polygenic phenotype impacted by variation throughout the genome, influenced by epistasis.

While the RFLP analyses demonstrated varied and complex recombination patterns, the resolution was not sufficient to allow confident identification of candidate genes and epistatic relationships influencing the observed phenotypic variation. We had previously reported that mixing of gH and gO alleles can epistatically impact the efficiency of fusion regulated by gH/gL/gO binding to the receptor PDGFRα and Ab neutralization on conserved anti-gH epitopes (41, 42). The short intergenic region between UL74 (gO) and UL75 (gH) shows relatively low LD (20) suggesting that crossovers can occur and this offers a potential explanation for differences among recombinants in specific infectivity and inhibition of spread by nAb. Among the recombinants, only E7 had a RFLP pattern consistent with a recombination mixing the TB and ME alleles of gH and gO since the UL74 matched TB and the UL82 matched ME. However, given the thousands to tens of thousands of Kbp between RFLP loci, crossover points could not be identified and *de novo* acquired point mutations could not be excluded. Moreover, since all three phenotypes analyzed are likely to be complex and polygenic, it may be that isolates can present with similar phenotypes for mechanistically distinct reasons. For example, one isolate may have a high specific infectivity principally because of efficient attachment and fusion, whereas another isolate with comparable infectivity has less efficient attachment and fusion but this is compensated by more efficient nuclear docking and uncoating and yet another because of subversion of the cellular antiviral state mechanisms. Genetic correlates associated with each of these scenarios would be expected to be different. Identification of strong candidate genes and epistatic relationships will require analysis and comprehensive sequence analyses of these recombinants and more from the original TB-ME coinfection.

The regions of the HCMV genome between the allelic genes show strong sequence conservation and signatures of recombination (20, 21). Questions remain as to whether these recombination signatures represent rare events accumulated over the evolutionary history of HCMV or an ongoing, dynamic process influencing HCMV pathogenesis and intervention successes and failures. Our results suggest that recombination may be a common outcome of coinfection of a single cell by 2 genetically distinct HCMV variants. This is consistent with the model of recombination-dependent replication that posits efficient DNA replication of herpesviruses requires recombination to cope with single and double stranded breaks (55). Given the dramatic phenotypic differences among the few strains and BAC clones that have been studied in the laboratory, and the allelic diversity among HCMV sequences in the database, the phenotypic variation of HCMV *in vivo* might be dramatically more vast and *in vitro* characterized phenotypes considered artifactual may be more physiologically relevant than currently appreciated Our approach of generating TB-ME recombinants is a step towards building *in vitro* models that may begin to more fully reflect the *in vivo* genetic diversity of HCMV.

## MATERIALS AND METHODS

### Cell lines

Primary human foreskin fibroblast cells (HFF; Thermo Fisher Scientific), and tetracycline repressor protein (TetR)-expressing HFF (HFFtet; (37)) were cultured in Dulbecco’s modified Eagle’s medium (DMEM, Sigma) supplemented with a mixture of 5% heat-inactivated fetal bovine serum (FBS, Rocky Mountain Biologicals, Missoula, MT, USA) and 5% Fetalgro® (Rocky Mountain Biologicals, Missoula, MT, USA), penicillin-streptomycin, gentamicin, and amphotericin B.

### Human cytomegaloviruses (HCMV)

BAC clone TB40/e-BAC4 (TB) was provided by Christian Sinzger (University of Ulm, Germany) (56). BAC clone Merlin (pAL1393), which contains tetracycline operator sequences within the transcriptional promotor of UL131, was provided by Richard Stanton (Cardiff University, United Kingdom) (31). BAC clones were modified to express green fluorescent protein (GFP) or the monomeric red fluorescent protein, mCherry (mC) with *en passant* recombineering replacing the US11 gene with the eGFP or mC gene (37). Infectious HCMV was recovered by electroporation of BAC DNA into MRC5 cells (33), which were then co-cultured with either HFF cells (TB) or HFFtet cells (ME). Supernatant virus was harvested and infectious units (IU) determined by flow cytometry (37). ME stocks propagated in HFFtet cells were denoted “MT”

### TB-ME recombinant isolates

Six confluent 150x20 mm tissue culture plates of HFFtet cells were inoculated with 7.6×10^6^ IUs (MOI = 0.25 IU/cell), combined of TBus11mCherry (TB_mC) and MTus11GFP (MT_GFP), diluted to 10 ml total volume per plate and incubated at 37°C for 4 hours. Inoculum was aspirated and replaced with DMEM+2% FBS The infections were incubated for 2 days before the cells were lifted with 0.5x trypsin and pelleted at 500g for 5 minutes. Half of the cells were resuspended in 1 mL sorting buffer (F-12/DMEM; Sigma +10% FBS) and the other half resuspended in 1 mL DMEM+2% FBS and frozen at −80°C. F-12/DMEM was used as the sorting buffer because it is HEPES buffered to maintain pH in ambient CO_2_ during sorting, approximately 3 hours. Cells were sorted using a BD Aria Fusion flow activated cell sorter (FACS). Cells were gated from debris using forward scatter-area and side scatter-area, and single cells were gated using forward scatter-width and forward scatter-height. GFP+ and mC+ gates were drawn using non-tagged virus-infected cells as negative controls. Approximately 2.5×10^6^ dual GFP+/mC+ were collected, representing about 40% of the total infected cells, and plated on a single 100x20 mm tissue culture plate. The survival rate was about 75% after three days. At 8 days post sorting (10 days post infection), supernatant was harvested and frozen at −80°C in 1 ml aliquots. IUs were determined as above, mC: 6×10^6^ IU/mL, GFP: 1×10^6^ IU/mL

To isolate individual progeny viruses, two 100x20 mm plates of confluent HFFtet cells were inoculated with the supernatant (0.4 μM filtered to remove cells, cellular debris and viral aggregates) in 5 mL/plate total volume (MOI of 0.009, and 0.18 IU/cell) for 4 hours at 37°C and the inoculum replaced with 10 ml of DMEM+2%FBS. Two MOI were used to decrease the probability of coinfections while maintaining enough infected cells for FACS collection. At three days post inoculation the cells were lifted with 0.5x trypsin from both and resuspended together in 1 ml sorting buffer. Simultaneously, twelve 96-well plates were seeded with HFFtet cells at a density that would results in about 125% confluency when adhered (planning for an average 75% survival rate). The infected cells were then sorted, using stringent gating for only GFP+ or only mC+ cells and one infected cell was deposited per well of the 96-well plates (6 plates of GFP+ cells and 6 plates of mC+ cells). Beginning at day 10 post sorting, wells were examined by fluorescence microscopy for focus formation. Wells with no foci or with both GFP and mC were discarded. Wells with only GFP or mC were harvested by scraping cells into the supernatant with a cut pipet tip and then cells and supernatant together were stored at −80°C. To amplify an isolate, the entire sample (supernatant and cells) was plated on one well of a 48-well dish along with fresh HFFtet cells at a density to achieve confluence. Most isolates were expanded after 10 days, although some slower spreading isolates were allowed to spread for an additional 10 days prior to expansion. In attempt to limit selection of cell-free or cell-associated viruses, expansion involved collecting the culture supernatant and the intact cells together and transferring to a 6-well dish along with fresh HFFtet. For experiments, high MOI “burst” supernatant stocks were generated on HFF (non TetR-expressing) as before (25)

### Restriction fragment length polymorphism (RFLP) genotyping

DNA was extracted from infected cells using PureLink genomic DNA minikit (Thermo Scientific) and PCR amplified with primers listed in Table 2. PCR amplicons were then digested with indicated restriction enzymes (NEB BioLabs) and analyzed on 1% agarose gels. Purified BAC DNA was used for experiments involving titration of parental TB and ME as PCR input.

### Immunoblotting

Supernatant virions were analyzed by immunoblotting as before (40). Briefly, virions were solubilized in 2% SDS–20 mM Tris-buffered saline (TBS) (pH 6.8) and separated on 4-20% gradient SDS-PAGE (BioRad) under nonreducing conditions and transferred to polyvinylidene difluoride (PVDF) membranes (BioRad) in 10 mM NaHCO_3_ and 3 mM Na_2_CO_3_ (pH 9.9), 20% methanol. Transferred proteins were detected with rabbit anti-gL (57) and anti-rabbit horseradish peroxidase (HRP) conjugated secondary antibody (Sigma). Following exposure to Pierce ECL Western blotting substrate (Sigma), chemiluminescence was detected using a Bio-Rad ChemiDoc MP imaging system.

### Quantitative PCR

Viral genomes were determined as described previously (25) Briefly, cell-free HCMV stocks were treated with DNase I before extraction of viral genomic DNA (PureLink viral RNA/DNA minikit; Life Technologies/Thermo Fisher Scientific). Primers specific for sequences within UL83 were used with the MyiQ real-time PCR detection system (Bio-Rad).

### Virus Spread

As a modification of the method described by Schultz (37), HFF cells were seeded in two 6-well tissue culture plates, per sample, at about 200,000 cells per well and allowed to grow to confluence. At 3 days post-confluence, cells were inoculated with about 1,000 IU/well of virus stock in 1 mL DMEM + 2% FBS and incubated at 37^0^C for 4 hours. Inoculum was then replaced with 3 mL per well of DMEM+2%FBS or 3 mL DMEM+2%FBS supplemented with 50 μg/mL anti-gH 14-4b monoclonal antibody. One 6-well plate was harvested at 3 days post infection by trypsinizing and fixing cells in 4% formaldehyde in PBS. The second plate was harvested at 12 days post infection in the same manner. Fixed cells were pelleted, washed, and re-suspended in 1 mL PBS. Infected cells were quantified by flow cytometry.

## DATA AVAILABILITY

All data supporting the findings of this study are presented within the article. Further details on the procedure of all experiments can be obtained from the corresponding author.

## ACKNOWLEDGMENTS

We are grateful to Bill Britt, Christian Sinzger, and Richard Stanton, for generously supplying HCMV BAC clones, antibodies, and cell lines, as indicated in Materials and Methods. Additionally, we are grateful to the University of Montana (UM) Center for Biomolecular Structure and Dynamics (CBSD) and Center for Environmental Health Sciences (CEHS) for expert advice and instrumentation in purification of monoclonal antibodies and flow cytometry.

This work was supported by a grant from the National Institutes of Health (NIH) to B.J.R (R01AI097274), a NIH CoBRE award to the UM CBSD (P30GM140963) and a NIH S10 equipment grant to the UM CHES (S10OD025019-01).

Experiments were designed by C.S.P., B.J.R., and J.-M.L. and performed by C.S.P, I.T.B., I.G. and J.-M.L., and the manuscript was prepared by C.S.P and B.J.R.

